# “Seipin mediates Perilipin-1 recruitment to lipid droplets to preserve human adipocyte identity”

**DOI:** 10.1101/2025.11.09.687445

**Authors:** Denise Zhong, Ioanna Stavrakaki, Anand Desai, Tiffany DeSouza, Sabine Linskey, Shannon Joyce, Courtney Hatton, Danna Garcia, Reyan Kassam, Gregory Hendricks, Keith Reddig, Guangping Gao, Jun Xie, Maria Paz Gonzalez-Perez, Andre F. C. Vieira, Caroline L. Chidley, John A. Haley, Silvia Corvera

## Abstract

Congenital generalized lipodystrophy type 2 (CGL2) is caused by mutations in the *BSCL2* gene, which encodes the protein seipin. However, how seipin loss causes adipose tissue failure remains unclear. Using human adipocyte progenitor cells capable of robust differentiation in vitro and in vivo, we reveal two unexpected findings that redefine CGL2 pathogenesis. First, seipin is dispensable for lipid droplet biogenesis but essential for recruiting the major adipocyte scaffold protein Perilipin 1 (PLIN1) to the lipid droplet surface. Second, we discover that the integrity of the lipid droplet serves as an organelle to nucleus quality control checkpoint enforcing adipocyte identity. Without seipin-dependent PLIN1 recruitment, adipocytes exhibit enhanced lipolysis and ceramide accumulation, triggering an unexpected cellular response of de-differentiation into a progenitor-like state. From this de-differentiated state, cells can undergo additional cycles of differentiation and de-differentiation upon repeated adipogenic stimuli. However, some cells escape de-differentiation, instead forming a single large droplet and displaying severe cellular structural abnormalities. Consistent with this model, we find functional adipose tissue can form in vivo from seipin-deficient cells, yet ultimately fails. These findings resolve conflicting models of CGL2 pathogenesis by reframing seipin as a regulator of PLIN1 recruitment, rather than droplet formation per se, and reveal the fundamental role of lipid droplet integrity in the development of functional human adipocytes.

---

Congenital generalized lipodystrophy type 2 (CGL2) is a rare and severe condition caused by loss-of-function mutations in *BSCL2*, the gene encoding Seipin. Individuals with CGL2 lose metabolically active and mechanical adipose tissue and develop early-onset metabolic complications, including insulin resistance, hepatic steatosis, pancreatitis, and premature mortality [1, 2]. Mouse models in which *Bscl2* deficiency is restricted to adipocytes recapitulate much of the human phenotype, suggesting that adipocytes are the cell type responsible for the CGL2 metabolic phenotype [3]. The cellular mechanisms by which loss of Seipin disrupts human adipocyte function remain poorly understood.

A major obstacle to progress has been the inability to directly study human adipocytes from affected individuals. The near-complete absence of adipose tissue in these patients precludes isolation of the disease-relevant cell type. As a result, most studies have relied on indirect or heterologous models— such as fibroblasts, lymphoblasts, or engineered non-adipose cell lines—that do not undergo adipogenesis or recapitulate the biology of human adipocytes. Similarly, animal models and studies in yeast or Drosophila have been critical for identifying conserved aspects of seipin function [4–6], but they do not fully capture the complexity of human adipocyte function or adipose tissue development.

To address these limitations, we developed a model based on the expansion of mesenchymal progenitor cells derived from human adipose tissue samples [7, 8]. These cells can be expanded extensively while maintaining adipogenic potential, are amenable to CRISPR-Cas9 editing [9, 10], and generate functional adipose tissue when transplanted into immunodeficient mice. Unlike previous models, this system recapitulates the full trajectory of human adipogenesis—from progenitor to mature adipocyte in vivo—providing direct access to the cell type and developmental processes that are disrupted in CGL2.

Using this approach, we find that loss of seipin prevents recruitment of the major droplet scaffold protein PLIN1 onto the emerging lipid droplet. This failure is accompanied by abnormal lipid metabolism and triggers a collapse of adipocyte identity, with many seipin-deficient cells reverting to a progenitor-like state. These findings provide direct evidence that seipin is required to establish and maintain the functional architecture of the adipocyte lipid droplet during human adipogenesis, and reveals that lipid droplet acts as a gatekeeper of the adipocyte’s functional identity. The results also highlight the utility of this human-derived platform for uncovering disease mechanisms in rare adipose disorders.

## RESULTS

To determine the effects of *BSCL2* deficiency, we used multipotent mesenchymal progenitor cells that were previously derived from fragments of surgically discarded human abdominal subcutaneous adipose tissue. The cells were electroporated with Cas-9 protein and sgRNA directed to the *BSCl2* gene, by methods previously described [7, 10, 11]. To control for the effects of electroporation and excision-induced DNA damage, we targeted a gene that is not expressed in these progenitor cells or in human adipose tissue (*OR2S2*), which will be referred from here on as control (CTRL). The efficiency of excision was found to be >95%, as assessed by Sanger sequencing (Figure 1A). We then examined the differentiation trajectory of control and BSCL2-KO cells by phase microscopy. Contrary to our expectations, we observed abundant lipid droplets appearing in response to adipogenic induction in both control and BSCL2-KO cells (Figure 1B). However, 7-9 days after induction, we observed a striking pattern in which some cells formed a large unilocular droplet, but most other cells showed smaller and sparser lipid droplets. We quantified the distribution of lipid droplet sizes across multiple fields and time points (Figure 1C). In control cells, the size of lipid droplets progressively increases, as seen by the continuous shift to the right of the distribution curve. In contrast, in BSCL2-KO cells the shift to the right stalled after day 9 and this was accompanied by the emergence of a population of abnormally large droplets. Side-by-side comparison of control and BSCL2-KO cells at each time point (Figure 1D), shows the appearance of very large droplets in BSCL2-KO cells at day 9, and a leftward shift in the distribution of most droplets between days 9 and 14. These results reveal that, rather than impaired droplet formation per-se, seipin deficiency causes altered lipid droplet dynamics, with heterogeneous effects across cells. To rule out the possibility that lipid droplets were forming in cells that had escaped CRISPR editing, we confirmed by immunostaining that the *BSCL2* protein (seipin) was undetectable in cells containing lipid droplets (Figure 1E).

**Figure 1.**
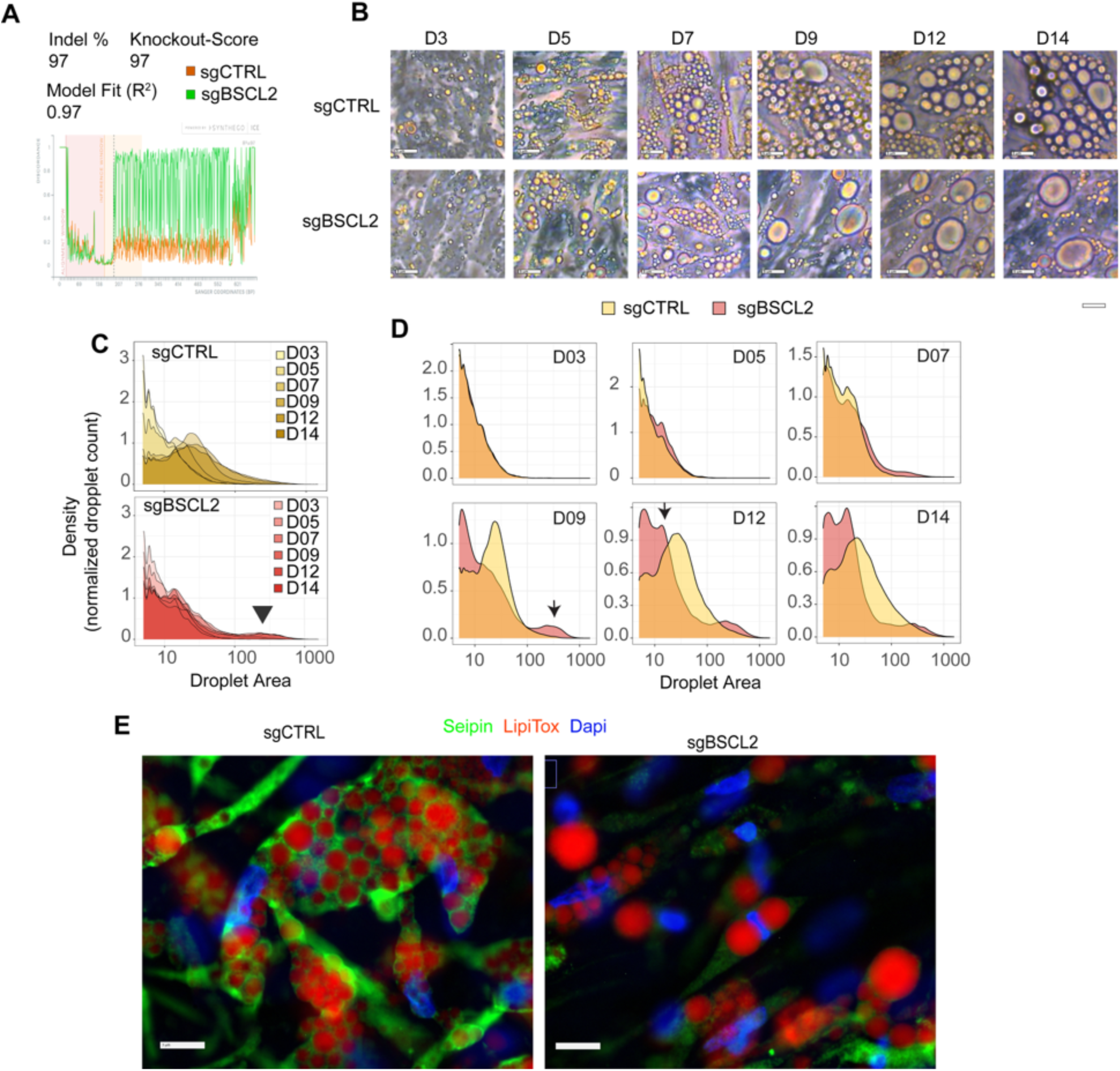
Altered lipid droplet dynamics in response to seipin deficiency. **A.** TIDE analysis of the BSCL2 gene locus in cells electroporated with Cas-9 and sgRNAs to a non-expressed gene (sgCTRL) or to the BSCL2 gene (sgBSCL2). **B.** Phase images of control or BSCL2-KO cells in which adipogenic induction was initiated at day 0. **C.** Density plots of lipid droplet distributions from day 3 (D3) to day 14 (D14) in control (top panel) and BSCL2-KO (bottom panel) cells. **D.** Density plots of lipid droplet distributions directly comparing control and BSCL2-KO cells at the indicated days after induction of differentiation. **E.** Immunostaining of control (left panel) and BSCL2-KO cells (right panel) with antibody to seipin (green), LipiTox (red), and Dapi (blue). Bars = 5µm.

PLIN1 is the major protein scaffold on adipocyte lipid droplets [12, 13]. To examine its potential role in mediating altered lipid droplet dynamics in seipin-deficient cells, we analyzed its localization by immunostaining during the differentiation trajectory. Strikingly, we found PLIN1 localization to the lipid droplet surface was compromised from the earliest stages of droplet formation in BSCL2-KO cells. At day 3 of differentiation, PLIN1 could be seen surrounding multiple small lipid droplets in both control and BSCL2-KO cells (Figure 2A, left panels). However, while in control cells PLIN1 surrounded the entire droplet (Figure 2A, top left panels), in BSCL2-KO cells PLIN1 formed patches close to the droplet but did not fully surround it. Moreover, we observed small speckles of PLIN1 in BSCL2-KO cells that were absent from control cells (Figure 2A, bottom left panels). As droplets enlarged over time, the defect in PLIN1 recruitment became more apparent (Figure 2A, right panels). Control cells accumulated many droplets, all of which were surrounded by PLIN1 (Figure 2A, top right panels). In contrast, BSCL2-KO droplets lacked PLIN1, which was instead distributed in a reticular interconnected network resembling the endoplasmic reticulum (Figure 2A, bottom right panels). These results indicate that seipin deficiency prevents the recruitment of PLIN1 from the endoplasmic reticulum to emerging lipid droplets.

**Figure 2.**
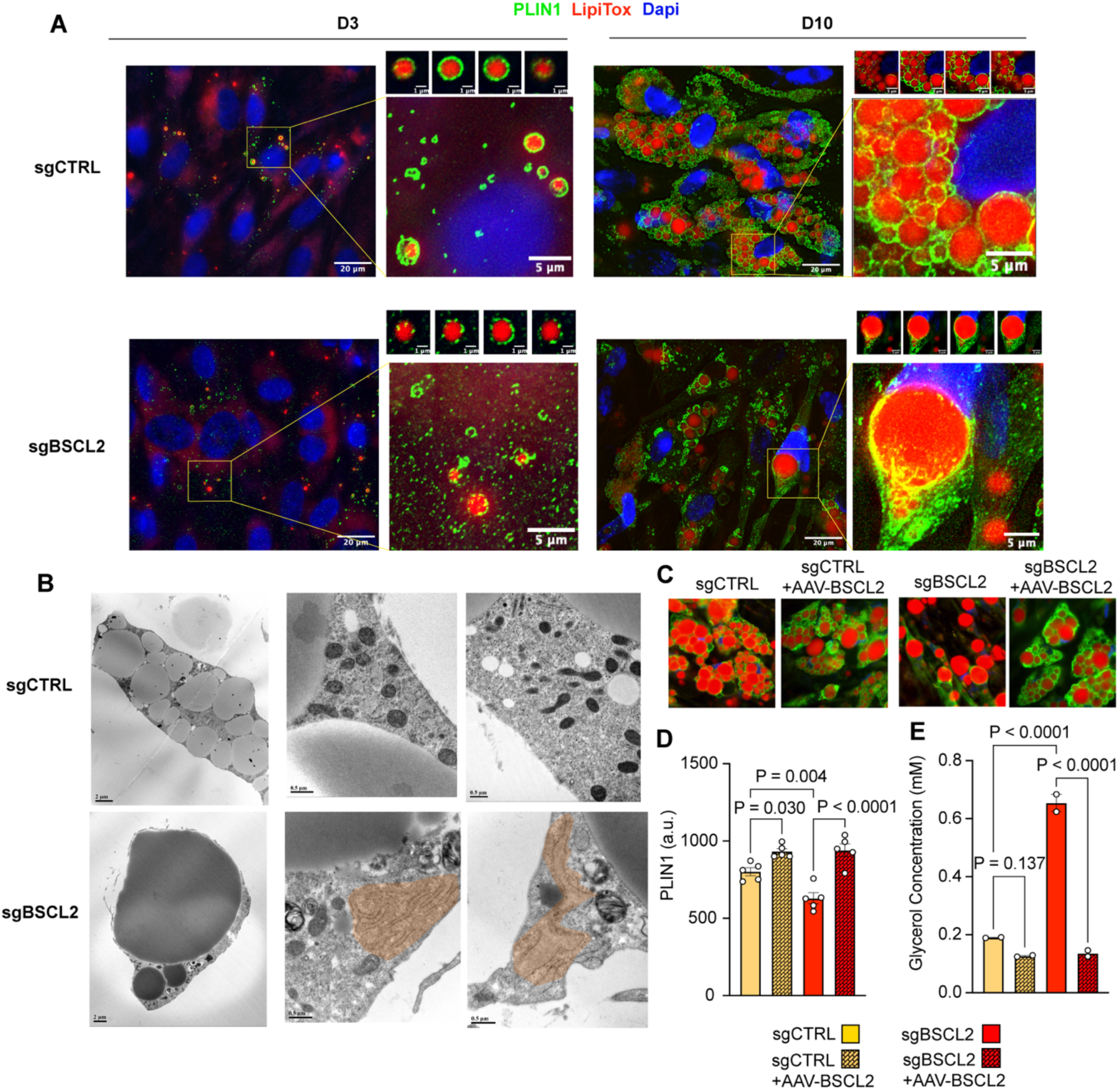
Seipin controls the recruitment of PLIN1 to lipid droplets. **A.** Immunostaining of control (sgCTRL) and BSCL2-KO cells (sgBSCL2) with antibody to PLIN1 (green), LipiTox (red) and Dapi (blue) at day 3 (left panels) and day 10 (right panels) after adipogenic induction. Wide-field images were captured on an Axiocam 705 camera using a Zeiss Plan-Apochromat 63x/1.40 Oil DIC M27 objctive a at 250 nm intervals, and deconvolved using 40 iterations of the regularized inverse filter algorithm through the DeconvolutionLab2 ImageJ plugin [43]. Maximal intensity projections of image stacks are shown. The areas marked by white rectangles are enlarged to the right, and individua optical sections of selected droplets are illustrated above the enlargements. **B.** Transmission electron micrographs of adipocytes at day 18 after induction of differentiation. Distended endoplasmic reticulum in BSCL2-KO cells is shaded in red. **C.** Control (sgCTRL) and BSCL2-KO cells (sgBSCL2) untransduced or transduced with AAV encoding human BSCL2 gene (AAV-BSCL2) stained with antibody to PLIN1 (green), LipiTox (red) and Dapi (blue). **D.** Quantification of PLIN1 signal relative to LipiTox in 5 independent fields. **E.** Quantification of glycerol accumulation in the media between days 9 and 10 in two independent wells. Statistical significance was calculated using one-way ANOVA and exact P-values are shown. Similar results were obtained in two additional experiments with cells from different donors.

To examine lipid droplet morphology at higher resolution, we conducted transmission electron microscopy (Figure 2B). After 10 days of differentiation, control adipocytes contained multiple droplets and well-defined cellular structures, while BSCL2-KO cells with large droplets displayed multilamellar vesicles, abnormal mitochondria, and a dilated endoplasmic reticulum. To verify that the observed structural abnormalities were indeed attributable to loss of seipin, we transduced control and BSCL2-KO cells with AAV-6 particles encoding human *BSCL2*/Seipin. Control cells contained normal lipid droplets surrounded by PLIN1, with higher PLIN1 intensity in cells transduced with AAV-BSCL2 upon quantification (Figure 2 C,D). BSCL2-KO cells displayed lower staining of PLIN1, which failed to surround lipid droplets, but BSCL2-KO cells transduced with AAV-BSCL2 displayed normal PLIN1 localization to multiple lipid droplets and were indistinguishable from control cells (Figure 2 C,D).

PLIN1 regulates lipolysis by preventing access of lipases to the lipid droplet triglyceride core. Consistent with this role, we find that the mislocalization of PLIN1 is accompanied by increased lipolysis, assessed by the concentration of glycerol in the cell culture media (Figure 2E). The reversal of PLIN1 mislocalization by AAV-BSCL2 was accompanied by complete mitigation of functional defects, with glycerol release becoming indistinguishable between control and AAV transduced BSCL2-KO cells (Figure 2E). These results confirm that recruitment of PLIN1 to lipid droplets is under the control of seipin. In the absence of seipin, PLIN1 fails to migrate to the lipid droplet, leading to structural and metabolic abnormalities.

PLIN1 controls basal lipolysis through modulation of the major triglyceride lipase, ATGL [14]. To determine whether increased lipolysis in seipin-deficient cells was attributable to ATGL, we generated a double knockout of seipin and ATGL (Figure 3A). ATGL knockout (87% efficiency) resulted in an increase in lipid droplet number and area at early stages of differentiation in control cells (Figure 3B), consistent with triglyceride accumulation resulting from inhibition of basal lipolysis. We find that the enhanced glycerol release in BSCL2-KO cells is mitigated by ATGL knockout but not fully restored to that seen in control cells (Figure 3C). Moreover, the abnormalities in lipid droplet distribution seen in BSCL2-KO cells were not reversed by ATGL depletion (Figure 3D). Density plots of lipid droplet sizes at day 10 of differentiation were similar when comparing control and ATGL-KO cells, and BSCL2-KO and double knockout cells. In contrast, clear differences are observed between control and BSCL2-KO cells, and between control cells and double knockout cells (Figure 3E). We conclude that the differences in lipid droplet dynamics and metabolic abnormalities in BSCL2-KO cells cannot be solely attributed to enhanced lipolysis stemming from ATGL activation.

**Figure 3.**
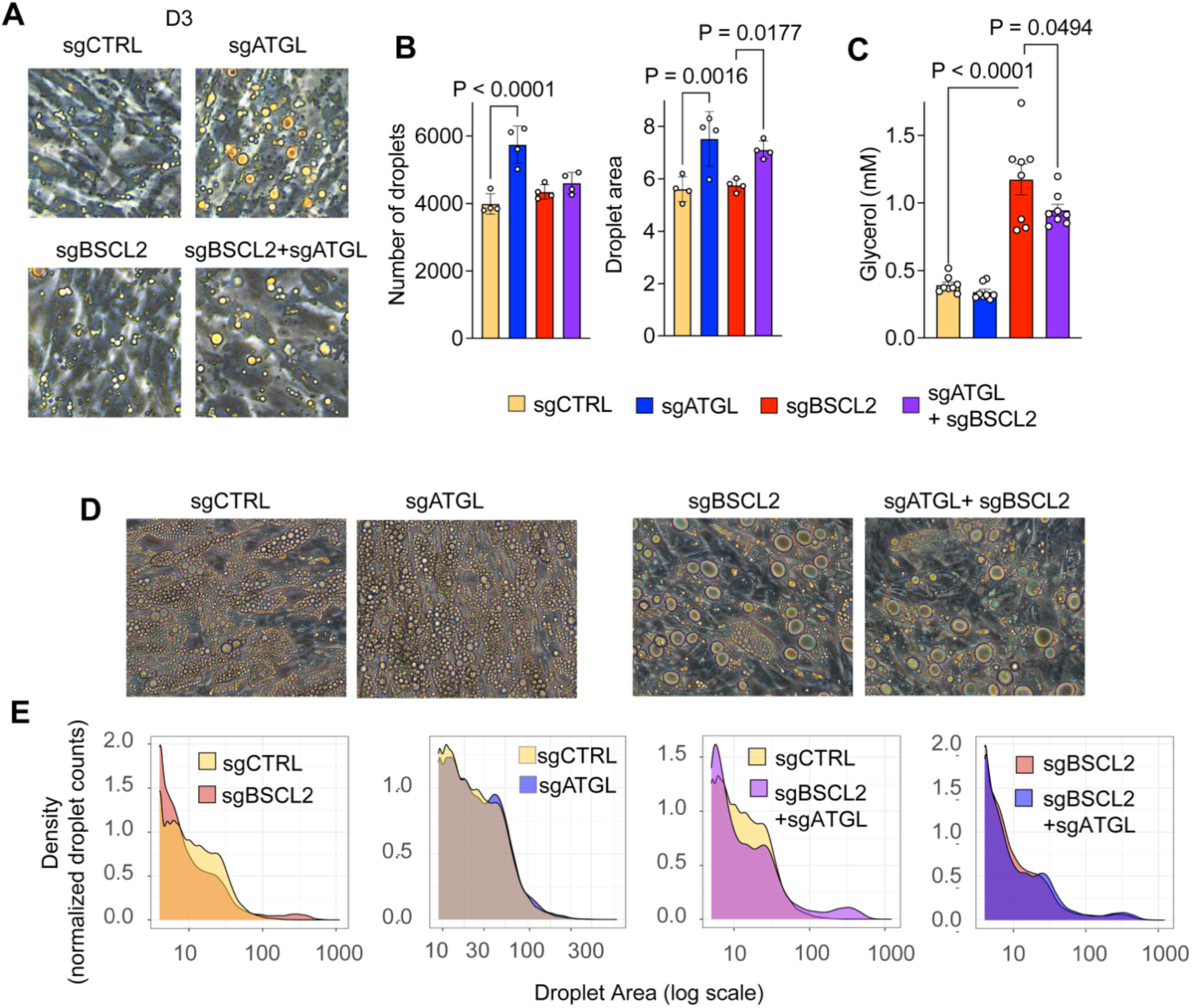
ATGL knockout fails to reverse the morphological and functional defects of BSCL2-KO cells. A. Cells electroporated with the indicated guide RNAs after day 3 of differentiation, showing emerging lipid droplets. B. Quantification of mean droplet number and mean area in fields from 4 independent cultures at day 3 of differentiation. Statistical significance was assessed using one-way ANOVA, and exact P-values are shown. C. Glycerol concentration in the media between days 9 and 10 of differentiation. D. Cells electroporated with the indicated guide RNAs after day 10 of differentiation. E. Density plots of lipid droplet size distributions comparing cells electroporated with indicated guides.

To examine whether seipin is required for the recruitment of other proteins, we conducted label-free protein mass spectrometry on lipid droplets isolated by flotation over sucrose gradients from cells gently disrupted using a Balch homogenizer (Figure 4A). We detected peptides corresponding to 7031 proteins, out of which 149 displayed 2-fold differences (Padj < 0.05) between control and BSCL2-KO cell lipid droplets (Figure 4B and Supplementary Table 1). 43 proteins were increased in droplets from BSCL2-KO cells compared to controls, the most salient being PLIN2 (27-fold increase), followed by CIDEC (8-fold increase). Notably, PLIN2 associates with nascent lipid droplets and is normally degraded once displaced PLIN1 [15, 16]. CIDEC mediates lipid droplet fusion, and its overexpression leads to accumulation of enlarged lipid droplets in adipocytes [17]. The overabundance of these proteins in lipid droplets from BSCL2-KO cells is consistent with failure of PLIN1 recruitment. Failure to recruit PLIN1 prevents displacement and subsequent degradation of PLIN2, leading to its accumulation, while excess CIDEC promotes droplet fusion and enlargement. The abundance of other PLINs was also increased in BSCL2-KO droplets (Figure 4C), consistent with the notion that PLIN1 normally limits the association of other PLINs originating from the endoplasmic reticulum or cytoplasm, and thereby loss of PLIN1 recruitment results in their accumulation on the droplet surface. To verify the results from mass spectrometry analysis, we stained control and BSCL2-KO cells with antibody to PLIN2. We find an increase in staining intensity in KO cells compared to controls starting at around day 5 of differentiation (Figure 4D,E). Moreover, while PLIN2 surrounded small lipid droplets and showed minimal colocalization with PLIN1 in control cells (Figure 4F, white arrows), it was highly co-localized with PLIN1 in a diffuse pattern in BSCL2-KO cells. These observations suggest that, in the absence of seipin, PLIN2 accumulates within both the endoplasmic reticulum and cytoplasm. Interestingly, PLIN1 abundance in the lipid droplet fraction did not appear decreased by mass spectrometry, contrary to expectations from imaging data. This apparent discrepancy is likely explained by retention of PLIN1 in ER fragments closely apposed to lipid droplets—structures visible by electron microscopy (Figure 4G)—which co-fractionate with droplets during isolation.

**Figure 4.**
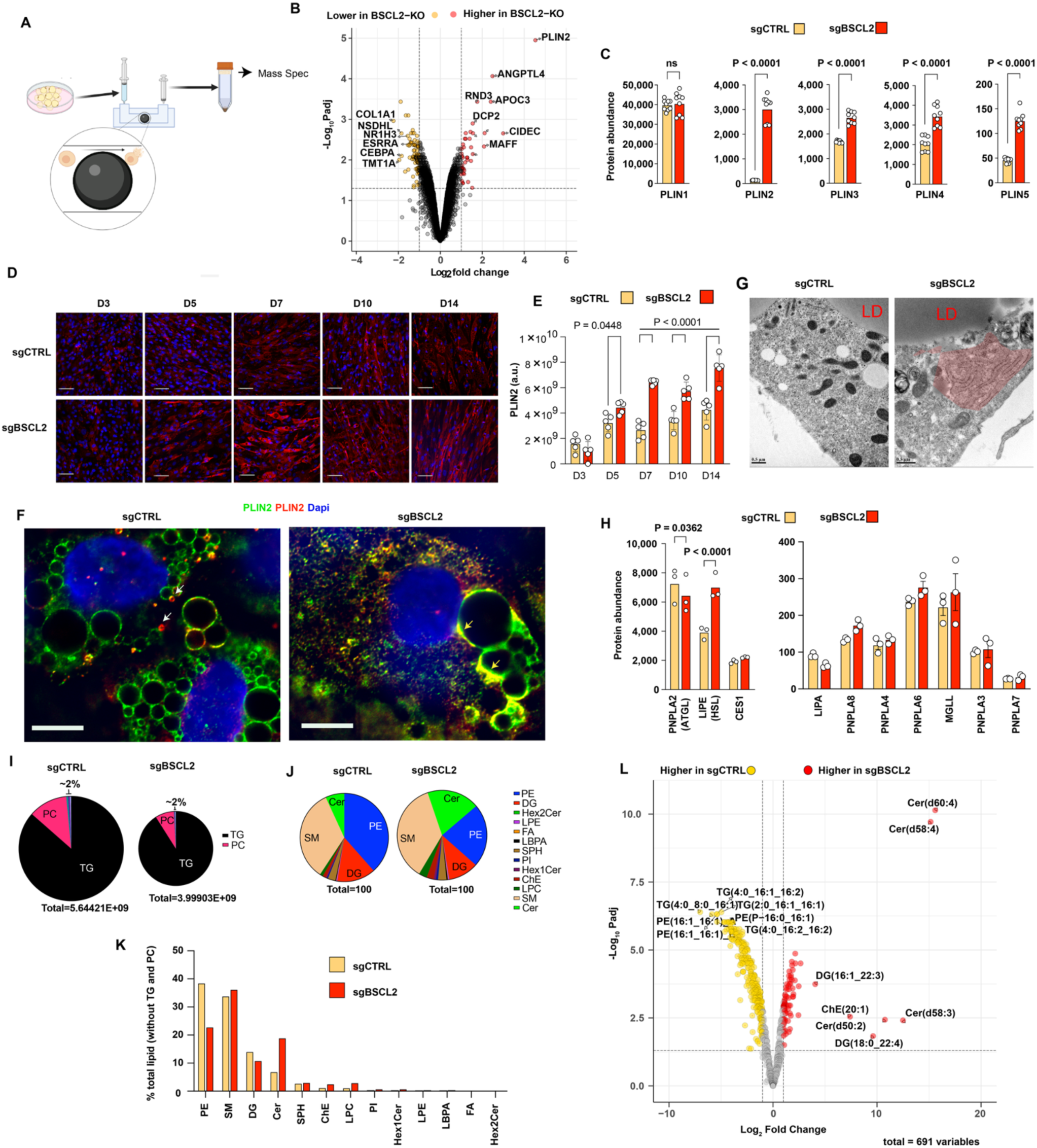
Proteomic and lipidomic analyses of lipid droplets. A. Schematic of procedure to obtain lipid droplets from cultured human adipocytes using sucrose gradient centrifugation after disruption using a Balch homogenizer. B. Volcano plot depicting changes in protein abundance between lipid droplets from control or BSCL2-KO adipocytes at day 10 of differentiation. The dataset comprised 18 LC-MS/MS runs across six biological conditions (sgCTRL 1,2,3 and sgBSCL2 1,2,3,) with three technical replicates per condition. The final analysis identified 182,724 precursors, corresponding to 125,671 unique peptides and 7,645 protein groups at 1% FDR. C. Abundance of indicated perilipins in the dataset. D. Immunofluorescence staining for PLIN2 in control and BSCL2-KO cells at indicated days of differentiation. Scale bars=100 µm. E. Quantification of PLIN2 signal intensity in 5 fields per time point. Statistical significance was estimated using one-way ANOVA and exact P-values are shown. F. Immunofluorescence staining of control and BSCL2-KO cells at day 5 of differentiation with antibodies to PLIN1 and PLIN2. Shown are individual optical sections after deconvolution. Small PLIN2 positive droplets seen in control cells are indicated with white arrows. Co-localization between PLIN1 and PLIN2 in diffuse regions around lipid droplets in BSCL2-KO cells is indicated with yellow arrows. Bars = 10 µm. G. Transmission electron micrographs of adipocytes, with expanded endoplasmic reticulum (shaded in red) in close apposition to the lipid droplet in BSCL2-KO cell. H. Abundance of indicated lipases in the dataset. Two graphs depict high abundance (left) and low abundance (right) species detected. I. Abundance of lipid species including triglycerides and phosphatidylcholine in lipid droplets in from control and BSCL2-KO cells. J,K Abundance of lipid species other than triglycerides and phosphatidylcholine in lipid droplets in from control and BSCL2-KO cells. L. Volcano plot depicting significant differences in specific lipid species in droplets from control and BSCL2-KO cells.

To gain insight on mechanisms underlying the increase in basal lipolysis seen in BSCL2-KO cells we examined the abundance of all detected lipases (Figure 4H). We find a small but significant decrease in ATGL, but a large increase in HSL, suggesting that increased diglyceride hydrolysis may underlie the observed increase in lipolysis. To gain a more comprehensive view of changes in lipid metabolism resulting from seipin deficiency and ensuing failure of PLIN1 recruitment, we conducted lipidomic analysis from the lipid droplet preparation. The total amount of lipid recovered, relative to protein yield, was lower in KO cells (Figure 4I), possibly reflecting contamination from the expanded endoplasmic reticulum, as hypothesized above. The major components corresponded to triglycerides, which reflect the core of lipid droplets, and phosphatidylcholine, their major surface phospholipid (Figure 4I). Other components included phosphatidylethanolamine, sphingomyelin, diglycerides and ceramides (Figure 4J,K). The most notable results from differential abundance analysis were a decrease in multiple triglyceride species and an increase in multiple ceramides in lipid droplets from BSCL2-KO cells (Figure 4L, Supplementary Table 2), which align with prior findings in adipose tissue of *Bscl2* deficient mice [3]. We detected decreased levels of triglycerides with short-chain fatty acids, which are difficult to detect but have been documented in mammals[18, 19]. Ceramides are associated with metabolic disease [20, 21], and notably ceramides with > 40 carbon chains have been identified linking lipotoxicity to endoplasmic reticulum stress in mouse skeletal muscle [22]. The mechanisms by which these unusual lipids are formed in seipin deficient cells, and their pathophysiological roles, warrant further investigation.

To gain insight on the cellular consequences of disrupted lipid metabolism, we conducted bulk RNASeq of control and BSCL2-KO cells generated from two separate donors. Hierarchical clustering of differentially expressed genes along the differentiation trajectory showed clustering by donor at days 0, 3 and 7 of differentiation, but clustering by knockout status at day 14, consistent with prior results indicating that major alterations due to seipin deficiency are not manifested until later stages of differentiation. Consistently, differential expression analysis showed no significant differences in gene expression at day 3 (not illustrated) or day 7 (Figure 5A) of differentiation. At day 14, however, numerous genes were decreased in BSCL2-KO cells from both donors (Figure 5B, Supplementary Table 3). Strikingly, enrichment analysis indicates that these genes map to adipocyte differentiation pathways, including PPARγ signaling, SREBP signaling, lipid biosynthesis and fatty acid metabolism.

**Figure 5.**
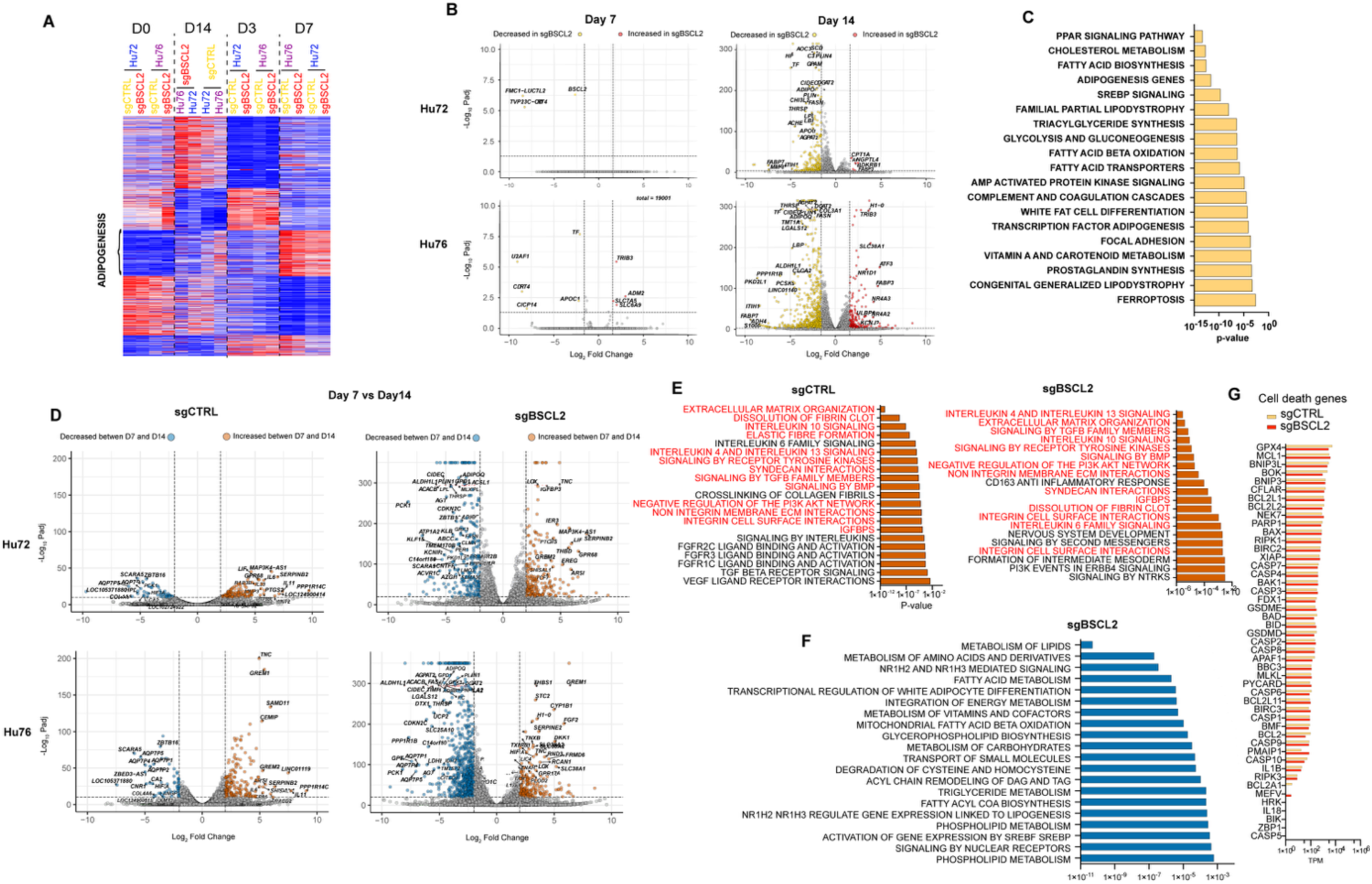
Seipin deficiency results in loss of adipocyte identity. **A.** Hierarchical clustering of differentially expressed genes across 32 RNASeq samples from two donors (Hu72 and Hu76) electroporated with either control (sgCTRL) or BSCL2-directed guides (sgBSCL2), with two biological replicates per condition, collected at days 0, 3, 7, and 14 of differentiation. Samples clustered by donor identity at days 0, 3, and 7, but by knockout status at day 14, indicating that transcriptional differences due to BSCL2 deficiency emerged during later stages of differentiation. **B.** Volcano plots depicting differences in expression between control and BSCL2-KO cells at days 7 and 14 from two donors, as indicated. C. Pathway enrichment analysis of genes decreased in BSCL2-KO (padj < 0.001, foldchange < 0.5), using WikiPathways. D. Volcano plots depicting differences in expression between days 7 and 14 of differentiation in control and BSCL2-KO cells from two donors, as indicated. E. Pathway enrichment analysis of genes increased between days 7 and 14 in control (sgCTRL) or BSCL2-KO (sgBSCL2) cells (padj < 0.001, foldchange < 0.5), using WikiPathways. Pathways shared between the two conditions are in red text. F. Pathway enrichment analysis of genes decreased between days 7 and 14 in BSCL2-KO cells (sgBSCL2). No enrichment was found when analyzing genes that decreased between days 7 and 14 in control cells. G. Mean TPM values for genes associated with cell death at day 14 of differentiation, comparing control and BSCL2-KO cells.

To better discern the changes in gene expression occurring between days 7 and 14 of differentiation, which is the period during which the effects of BSCL2 deficiency become manifest, we performed DESeq analysis between days 7 and 14 (Figure 5D, left panels, Supplementary Table 3). In control cells, 233 genes were increased (> 4x, Padj <0.001) but only 49 were decreased (< 0.25, P< 0.001) in cells from both donors. Genes that were increased mapped to pathways related to extracellular matrix organization and interactions, FGF, TGFb, VEGF, interleukin and tyrosine kinase signaling (Figure 5E). Genes that were decreased did not map to any pathway but interestingly included four genes encoding different aquaporins. In BSCL2-KO cells, 187 genes were increased, and many more genes (262) were decreased between days 7 and 14 in cells from both donors (Figure 5D, right panels). Notably, genes that were increased in BSCL2-KO cells mapped to similar pathways as those increased in CTRL cells (Figure 5E, red text), indicating that BSCL2-KO cells remain viable and continue to progress along a differentiation trajectory. Only 22 genes were increased between days 7 and 14 exclusively in BSCL2-KO cells, and included DPP4, a marker for adipocyte progenitor cells (Supplementary Table 3). In contrast, the 262 genes that were decreased in BSCL2-KO cells mapped specifically to lipid metabolism and adipocyte differentiation pathways. This specific decrease in adipogenic gene expression, despite continued upregulation of other pathways, suggests that adipogenic genes are selectively responsive to lipid droplet integrity, and that absence of seipin elicits a de-differentiation process. Importantly, we found no evidence of cell death, as assessed by expression of genes associated with major cell death pathways (Figure 5F), by staining for markers of these pathways (data not shown), or by presence of detached cells in the cultures.

The finding that lack of seipin elicits a de-differentiation process led us to examine whether pharmacological stimulation of the major regulator of adipose differentiation, PPARγ, might mitigate the BSCL2-KO phenotype. We treated cells with rosiglitazone (Rosi), a potent PPARγ agonist, starting at day 3 of differentiation, well before alterations become apparent in BSCL2-KO cells. We then acquired phase images to measure lipid droplet size distribution through the differentiation trajectory (Figure 6 A,B). We find that addition of Rosi increased the number of cells entering the adipocyte differentiation pathway, as evidenced by the larger number of droplets within all distribution ranges at all time points examined. The addition of Rosi did not rescue the lack of seipin; as seen in control cells, the number of droplets within all distribution ranges was increased, but neither the generation of very large droplets, clearly seen at day 9, nor the shift to smaller droplet sizes seen between days 9 and 12, were prevented by the PPARγ agonist. We also analyzed the distribution of PLIN1 in cells treated with Rosi. Despite displaying an increase in expression, PLIN1 remains predominantly in a diffuse distribution in Rosi-treated BSCL2-KO cells (Figure 6C). Functionally, both glycerol and free fatty acid concentrations in the media were higher in BSCL2-KO cells, both in the absence or presence or Rosi, indicating that lipolysis remains elevated. Moreover, whereas Rosi suppressed free fatty acid production in control cells, it failed to do so in BSCL2-KO cells, consistent with deficiencies in triglyceride synthesis evidenced by lipidomic analyses.

**Figure 6.**
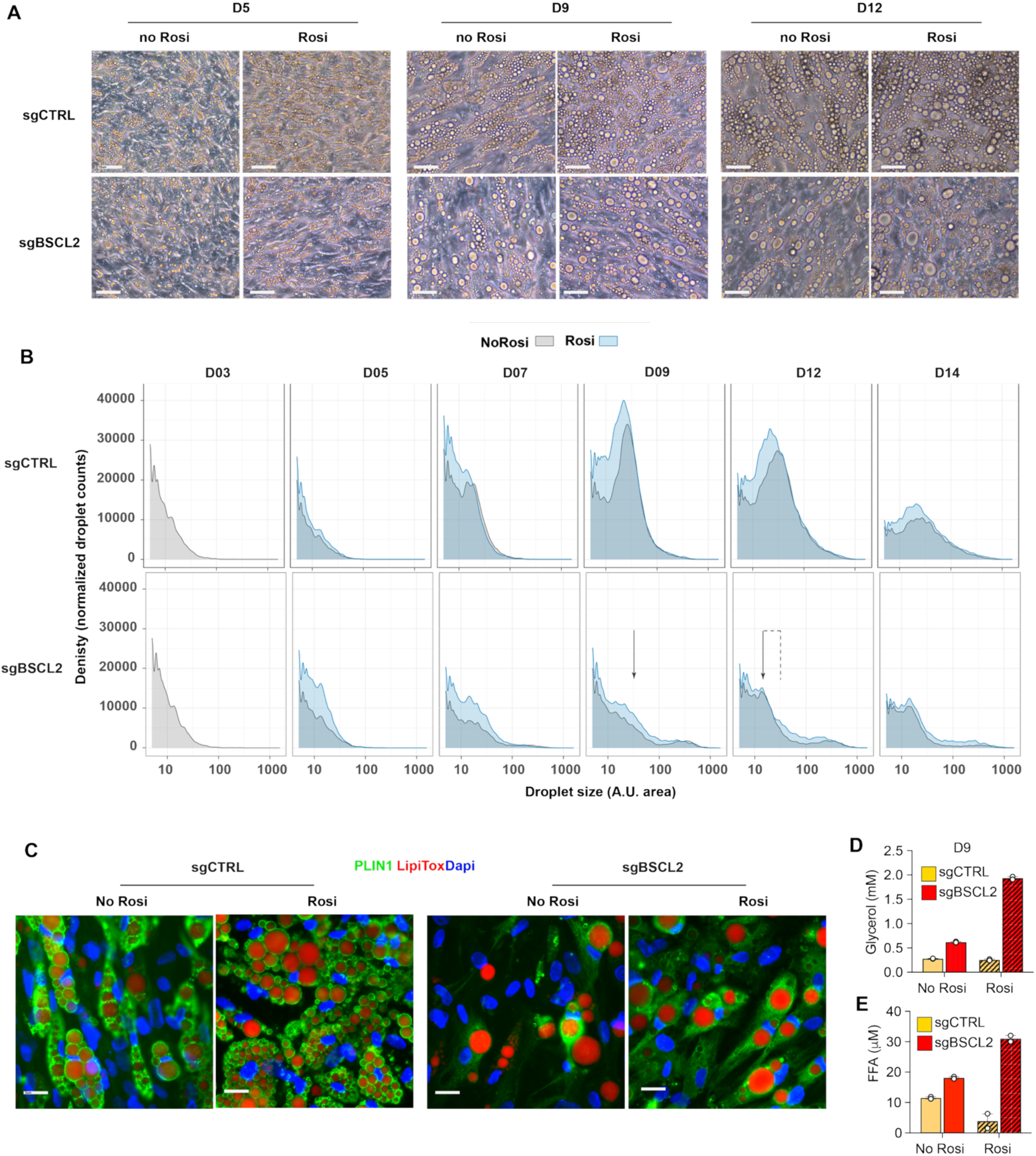
PPARγ activation does not restore the adipogenic program in seipin deficient cells. A. Images of cells cultured in the absence or presence of rosiglitazone starting at day 3 of differentiation, at days 5, 9 and 12. Scale bars=10 μm B. Density plots of lipid droplet sizes in control (upper panels) or BSCL2-KO cells (lower panels) treated without or with rosiglitazone at the indicated days. In control cells (sgCTRL) rosiglitazone increased the number of droplets, as seen by the higher peaks at all time points. An increase in lipid droplets is also seen in BSCL2-KO cells (sgBSCL2), but the leftward shift in droplet size is seen between day 9 and day 12 in BSCL2-KO cells is not prevented by rosiglitazone (arrows). C. Control or BSCL2-KO cells cultured in the absence or presence of rosiglitazone at day 9 after differentiation stained with an antibody to PLIN1 (green), LipiTox (red) and Dapi (blue). Scale bars=10 μm. D, E. Glycerol (D) and free fatty acid (µM) (E) concentrations in the media of control or BSCL2-KO cells cultured in the absence or presence of rosiglitazone at day 9 of differentiation. Results are derived from two independent wells.

To further explore the hypothesis that loss of seipin triggers dedifferentiation, we analyzed adiponectin secretion along the differentiation time course (Figure 7A). Culture media was replaced every 48 hours, so each measurement represents cumulative secretion during that interval. Adiponectin became detectable in the culture media at day 5 of differentiation and increased similarly in both control and BSCL2-KO cells through day 7. However, between days 7 and 9, adiponectin concentration began to decrease in BSCL2-KO cells compared to controls, and by days 9-11, secretion became nearly undetectable in BSCL2-KO cultures while remaining elevated in controls. To examine whether decreased adiponectin secretion was associated with cell death, we measured lactate dehydrogenase (LDH) activity in the culture media (Figure 7B). LDH activity was similar between control and BSCL2-KO cells throughout the time course, with only a transient elevation in BSCL2-KO cells between days 7 and 9 that subsequently normalized to control levels for the remainder of differentiation. These results indicate that the loss of adiponectin secretion is not due to cell death, consistent with the lack of cell death gene expression in BSCL2-KO cells observed by RNASeq (Figure 5G). Instead, the decline in adiponectin secretion, which coincides temporally with decreased expression of adipogenic genes, supports a model of dedifferentiation rather than cell death.

**Figure 7.**
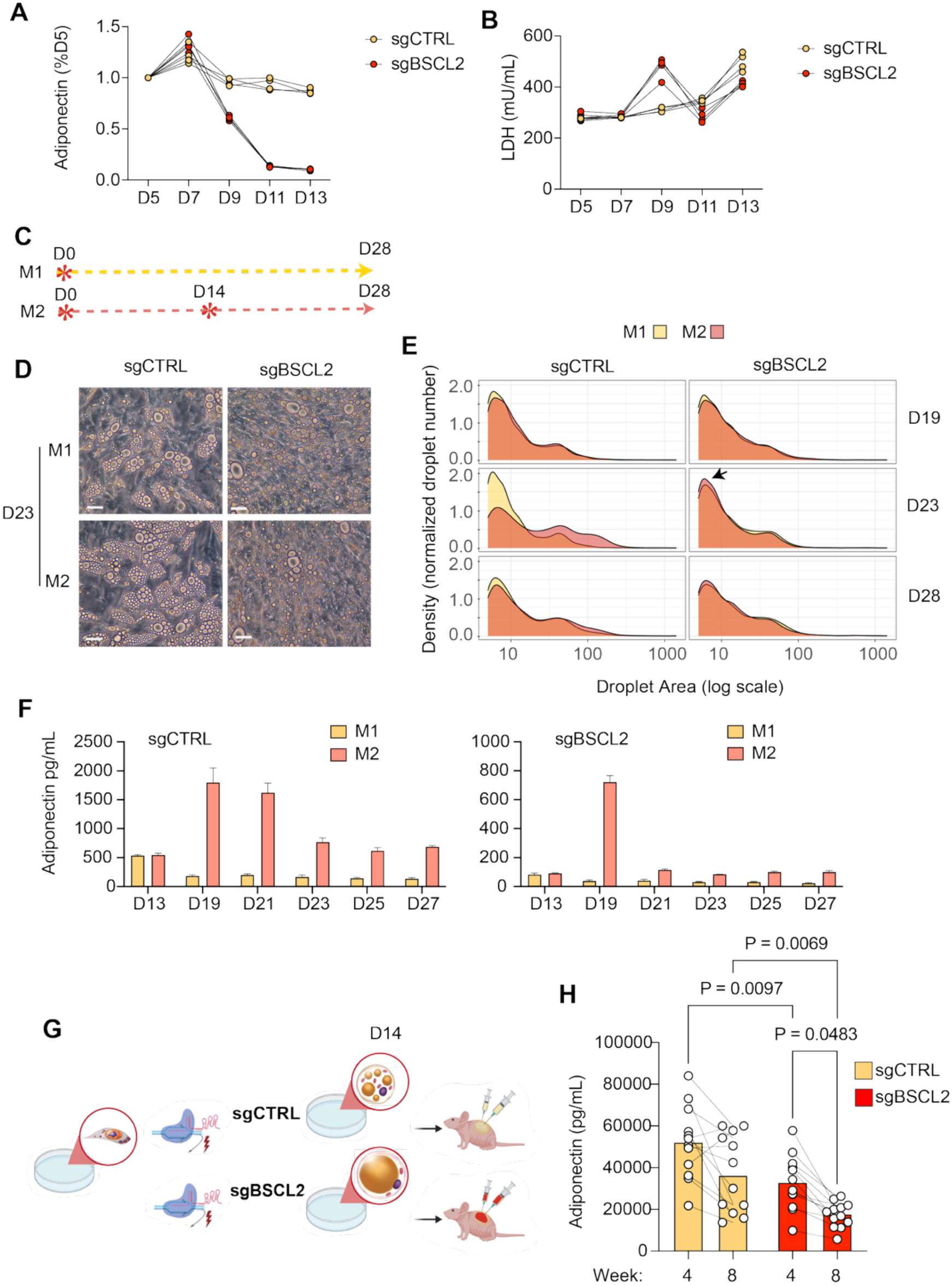
Evidence for re-differentiation following de-differentiation of seipin-deficient cells. **A,B.** Adiponectin concentration (A) and LDH activity (B) in the media of control (sgCTRL) or BSCL2-KO (sgBSCL2) cells throughout the differentiation trajectory. Shown are values from 4 independent wells. **C.** Schematic of experiment to test capacity of cells to undergo a second-round differentiation. At the indicated time point (stars), cells were induced with a mixture of insulin, dexamethasone and methyl isobutylxanthine. M1 depicts cells stimulated once, M2 depicts cells stimulated twice, at days 0 and 14. **D.** Images of control or BSCL2-KO M1 and M2 cultures at day 23. **E.** Density plots of cell size distributions in control (sgCTRL) and BSCL2-KO cells (sgBSCL2) comparing M1 and M2 at the days indicated on the right. **F.** Adiponectin concentration in the media of control (sgCTRL, left side plot) and BSCL2-KO cells (sgBSCL2, right side plot) comparing M1 and M2 at the days indicated. Arrow indicates an increase in number of small droplets in re-stimulated BSCL2-KO cells. **G.** Schematic for experiment to test capacity of BSCL2-KO cells to form adipose tissue in vivo. Human progenitor cells were electroporated with Cas9 protein and either control or BSCL2 guide RNAs were expanded in 15 cm dishes and differentiated for 14 days. Cells were trypsinized, resuspended in MatriGel, and implanted subcutaneously in the flanks of 6 male and 6 female NSG mice. **H.** Concentration of human adiponectin in the circulation of mice at weeks 4 and 8 following implantation. Each symbol represents an individual mouse. Statistical significance of the differences was estimated using one way ANOVA, and exact P-values are shown.

To determine whether cells that undergo de-differentiation might be capable of redifferentiation upon a second adipogenic stimulation, we exposed cells to differentiation cocktail twice, once at day 0, and another at day 14 (Figure 7C). Analysis of lipid droplet morphology indicated that control and BSCL2 cells responded differently to the second adipogenic stimulation (Figure 7D,E). In control cells we noted a shift to the right in the lipid droplet density distribution, consistent with an enlargement of pre-existing smaller droplets (Figure 7E, left panels). In contrast, there was a small increase in number of small droplets in cultures of BSCL2-KO cells (Figure 7D,E right panels, arrow). To determine whether the accumulation of droplets represents a second round of differentiation, we measured adiponectin in the culture media (Figure 7F). Both control and BSCL2-KO cells responded to the second adipogenic stimulation with increased adiponectin production. As was observed after a single stimulation (Figure 7A), adiponectin production by BSCL2-KO cells decreased faster than in control cells (Figure 7F, right panel). These results suggest that cells that undergo de-differentiation remain capable of renewed rounds of differentiation and de-differentiation.

To examine how this cellular behavior manifests in an in-vivo setting, we implanted control or BSCL2-KO cells into immunocompromised NSG mice (Figure 7G). To monitor human adipocyte development in the mice, we measured circulating adiponectin with an ELISA assay that specifically detects human and not mouse adiponectin (Figure 7H). After 4 weeks of implantation, we detected human adiponectin in the circulation of both male and female mice. After 8 weeks, a decrease in adiponectin production was seen, consistent with our prior published observations, which coincides with increased adipocyte size occurring in-vivo. Remarkably, BSCL2-KO cells were able to develop functional tissue, as evidenced by the concentration of adiponectin after 4 weeks, which was clearly detectable, albeit lower than that in mice implanted with control cells. However, the decline in adiponectin production after 8 weeks was more pronounced in mice implanted with BSCL2-KO cells. These results suggest that, in-vivo, cycles of differentiation and de-differentiation can support BSCL2-KO adipocyte function for a significant period.

## DISCUSSION

The most severe form of lipodystrophy is seen in humans with mutations in the *BSCL2* gene, encoding seipin. Seipin has been associated with lipid droplet assembly since it emerged as a candidate in a screen conducted in yeast to detect genes important for lipid droplet morphology [23]. Yeast seipin was localized to puncta on the endoplasmic reticulum of yeast, and its absence resulted in formation of irregular lipid droplets clustered alongside proliferated endoplasmic reticulum (ER). Very enlarged lipid droplets were also seen. Cryo-electron microscopy and structural modeling data of S. cerevisiae seipin revealed a decameric, cage-like structure with transmembrane segments interacting in two distinct, alternating conformations. In addition, data suggested that seipin attracts triglyceride monomers from the ER to sites of droplet formation [24]. These findings suggested a model in which seipin enables triacylglycerol phase separation, and subsequent lipid droplet growth and budding [6].

Despite its ubiquitous presence and function in many cell types, the dominant feature of *BSCL2* mutations in humans and mouse models is generalized lipodystrophy, with its severe associated metabolic disturbances. Thus, the function of seipin is especially relevant in adipocytes. Studies directed at understanding the mechanism by which loss of seipin results in loss of adipose tissue have identified multiple defects, including enhanced lipolysis attributed to uncontrolled PKA-mediated phosphorylation [25], altered regulation of lipin 1, a phosphatidic acid phosphatase [26], as well as impaired cytoskeletal remodeling [27]. These defects have been associated with defective adipocyte differentiation [28], attributed to failure of lipid droplet formation, although formation of very large droplets in seipin-deficient adipocytes has been consistently observed [3]. Our finding that seipin is required for the recruitment of PLIN1, an abundant lipid droplet scaffold protein specific to adipocytes and central to the control of lipolysis, conciliates many of these findings, PLIN1 was the first perilipin discovered [29], and was described as a major protein in adipocyte lipid droplets [13]. PLIN1 is specific for adipocytes, where it functions as a central regulator of lipid homeostasis. Four additional perilipins have been discovered subsequently, which are ubiquitously expressed [12]. The perilipins are recruited to lipid droplets by two general mechanisms: some are synthesized as monotopic integral membrane proteins at the ER and subsequently move to the lipid droplet surface using an ER to lipid droplet targeting pathway, while others can be recruited from the cytoplasm [30] [31]. PLIN1 is recruited from the ER, consistent with our observation that it accumulates in ER-like structures in the absence of seipin. In addition, our finding of a large increase in PLIN2 in seipin knockout cells can be attributed to its recruitment and stabilization on lipid droplets in the absence of PLIN1[32, Kaushik, 2016 #120].

PLIN1 also prevents access of other proteins to the lipid droplet, including lipases and lipase activators that target the triglyceride core [14, 33]. Thus, the absence of PLIN1 from lipid droplets elicits lipolysis, attributable to uncontrolled access of triglycerides to activated lipases. Consistent with a failure of PLIN1 recruitment, we and others have observed increased lipolysis upon knockdown of PLIN1 [9, 34] similar to that observed in seipin deficient cells (Figure 2E, Chen, 2012 #81}. Notably, increased lipolysis is not solely attributable to uncontrolled activation of the major PLIN1-regulated lipase ATGL, as a double knockout of seipin and ATGL also displayed enhanced lipolysis (Figure 3). This finding is consistent with deregulated activity of additional lipases, such as HSL, which was increased in lipid droplets from seipin deficient cells.

Additional support for a critical role of PLIN1 recruitment to adipocyte lipid droplets underlying CGL2 are the findings that mutant alleles of PLIN1 result in rare forms of congenital lipodystrophy [35], and autoantibodies to PLIN1 are associated with acquired generalized lipodystrophy [36–38] with similar manifestations as CGL2. Together, these findings support the loss of seipin-mediated PLIN1 recruitment to lipid droplets as a molecular pathogenic cause of CGL2.

A surprising finding in our study was that the abnormalities in lipid droplet function ensuing from loss of seipin-mediated PLIN1 recruitment to lipid dropkets trigger a program of de-differentiation. Our findings are consistent with earlier studies in which the initial induction of adipogenic transcription factors, and of key genes mediating triglyceride synthesis, including AGPAT2, lipin 1, and DGAT2, was preserved in cells lacking seipin. However, the expression of these critical factors was not sustained [28]. Our results indicate that more than 200 genes associated with adipogenesis are coordinately shut down within a precise window after start of differentiation. This is not accompanied by cell death, and indeed, other genes associated with progression towards a later stage of adipogenic differentiation are not affected. These results suggest that lipid droplets communicate with the nucleus to ensure appropriate adipocyte function along the differentiation trajectory.

The specific mechanisms by which progression to a mature adipocyte state is halted are not yet known, but we believe involve mechanisms other than PPARγ activation. We conclude this because activation of PPARγ by rosiglitazone was not able to suppress the leftward shift to a smaller droplet size characteristic of seipin deficient cells, nor was able to mitigate enhanced lipolysis. This failure was not due to lack of PPARγ or PPARγ responsiveness, as we noted a a strong effect of Rosi to increase the abundance of PLIN1 in cells containing large lipid droplets (Figure 6), which provides clear evidence that PPARγ activation was effective in these cells. Nevertheless, activation of PPARγ was able to increase the number of cells entering the adipogenic trajectory. Because early adipocyte development and function are not impaired by seipin deficiency, an increase in recruitment of progenitor cells to the adipogenic trajectory can explain the positive effects of pioglitazone and rosiglitazone to mitigate the metabolic consequences of seipin deficiency in mouse models [3, 39–41] and in humans with CGL2[40]. However, these effects depend on a continuous supply of progenitor cells, which may be depleted over the lifetime.

A salient finding in our lipidomic analysis on lipid droplets from seipin deficient cells was the increase in abundance of several ceramide species. Increase in ceramides was noted in adipose tissue of mice with a specific knockout of seipin in adipose tissue [3], but the nature of the cell type in which these lipid abnormalities occur, which could include inflammatory macrophages in the tissue, were not elucidated. Our data indicate that increased ceramides are produced in adipocytes in response to seipin deficiency and identify highly unusual species with very long chain fatty acids being upregulated in seipin deficient cells. These long-chain species are consistent with dysregulation of enzymes involved in elongation of fatty acids, and specific ceramide synthases.

It is notable that some cells escape the lipid droplet-nuclear checkpoint and continue to accumulate triglycerides, forming a single large droplet. While these adipocytes may represent specific adipocyte subtypes present in these cultures [8], the emergence of cell subsets with an enlarged droplet was first noted in original screens identifying yeast seipin, where approximately 30% of yeast cells display greatly enlarged droplets, while other retained small droplets in apposition to the ER. Thus, a response to abnormal lipid droplet architecture appears to be an evolutionarily conserved feature of cells but is not a sufficiently stringent mechanism as cells stochastically escape the checkpoint with relatively high frequency. Adipocytes with enlarged droplets display numerous abnormal features, including a distended ER, multilamellar structures, mitochondrial with abnormal morphology, and failure to secrete adiponectin (Figure 7).

Most cells, however, undergo de-differentiation and can complete additional cycles of differentiation and de-differentiation upon repeated adipogenic stimuli. This ability likely accounts for the successful formation of adipose tissue in mice from seipin-knockout cells shown in Figure 7. However, the stochastic loss of cells due to checkpoint failure eventually results in tissue loss and the metabolic manifestations of lipodystrophy. These findings suggest potential therapeutic approaches for CGL2, either by strengthening the checkpoint or by preventing maturation and preserving adipocytes in an immature yet functional state. Further studies in this experimental system will enable exploration of genes or drugs that elicit these responses.

## METHODS

### Cells

Banked progenitor cells previously obtained from de-identified fragments of human adipose tissue by methods previously described [7] were used in this study. In brief, fragments of adipose tissue are embedded in MatriGel (Corning, 356231), and cultured for 14 days to allow proliferation of progenitor cells within the tissue fragment. Cells are collected after dissolving the MatriGel with dispase and selected by plating in plastic culture dishes, expanded for two passages, and cryopreserved. For this study, cryopreserved cells were thawed and expanded in media supplemented with EGM-2 MV endothelial growth factors (Lonza, Basel, Switzerland; BulletKit, CC-3202), electroporated as described below, and plated in plastic culture dishes or coverslips as described in each experiment. Upon reaching 100% confluency, adipocyte differentiation was initiated by replacing the growth medium with differentiation medium composed of DMEM (GIBCO/Thermo Fisher, 11995-073) supplemented with 10% v/v fetal bovine serum (FBS, Genesee Scientific, 25-514), 500 µM 3-isobutyl-1-methylxanthine (Sigma, I5879), 1 µM dexamethasone (Sigma, D1756), and 1 µg/mL (170 nM) insulin (Sigma, I5500) (MDI cocktail). Differentiation was induced over a 72-hour period, after which the medium was replaced with DMEM (GIBCO/Thermo Fisher, 11995-073) supplemented with 10% v/v FBS (Genesee Scientific, 25-514). Culture medium was replenished every 48 hours throughout the differentiation period.

### CRISPR-Cas9 Ribonucleoprotein (RNP) Assembly and Electroporation

Ribonucleoprotein (RNP) complexes were assembled using SpyCas9 protein (PNA Bio) and synthetic single-guide RNAs (IDT and Synthego sgRNAs; **Table 1**). RNPs were prepared in Buffer R (Neon NxT Resuspension R buffer, ThermoFisher A54298-02) at final concentrations of 3 μM SpyCas9 and 4 μM sgRNA, maintaining a molar ratio of 4:3. Complexes were incubated at RT for 20 min prior to electroporation. For the Double Knock-Out experiment, RNP complex mixes were prepared using a 1:1:2 ratio of respective sgRNAs to SpyCas9 (2μM sgBSCL2, 2μM sgATGL, 4μM SpyCas9). Each sgRNA was first incubated with SpyCas9 individually, and the two resulting RNP complexes were then combined prior to electroporation.

**Table 1.**
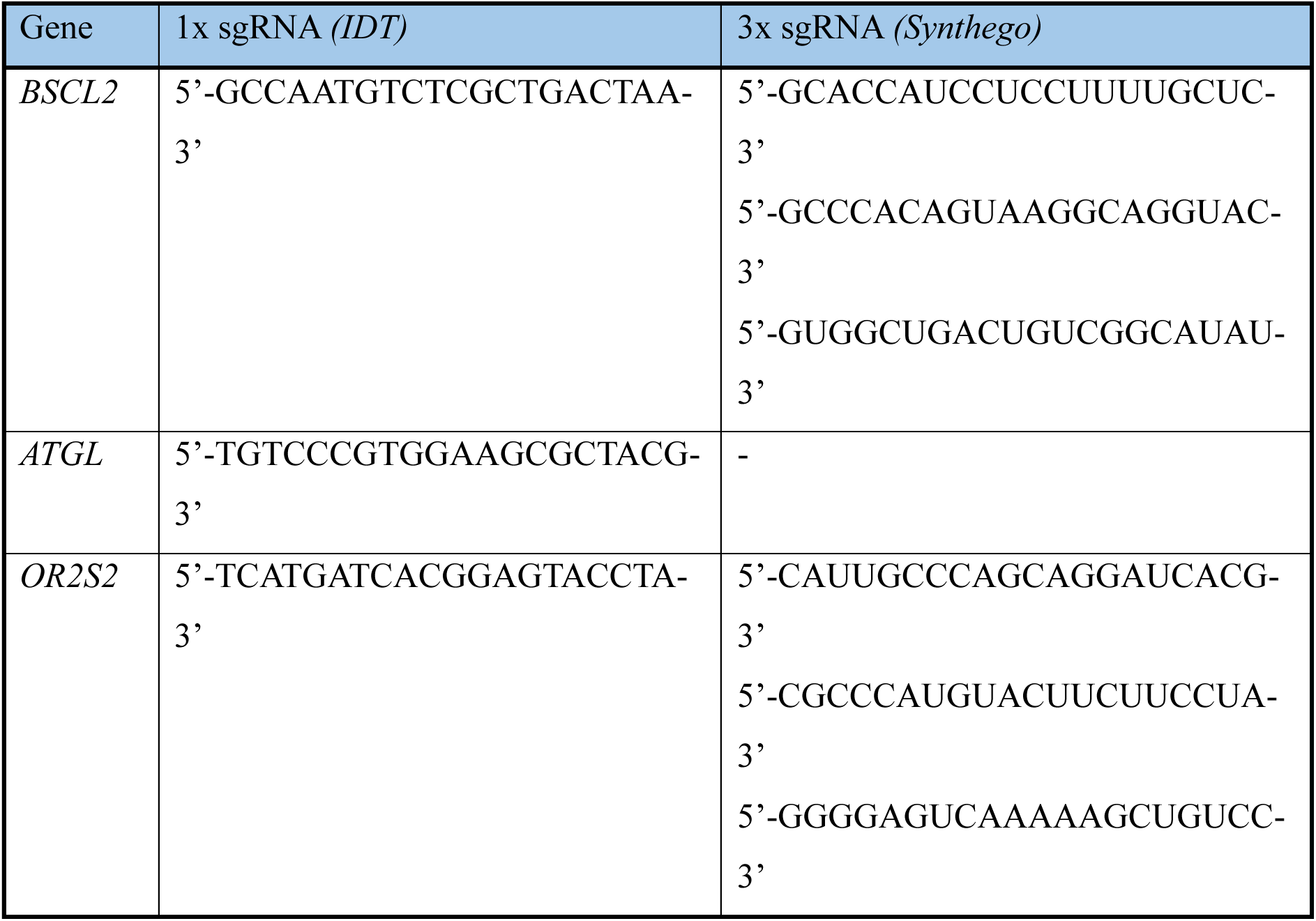
Single-guide RNA sequences used for gene modification using CRISPR-Cas9.

Cells were resuspended in Buffer R to a final concentration of 2-2.5 × 10⁶ cells per electroporation pulse (100μL cell-RNP mix/ pulse). The cell suspension was then mixed in a 1:1 ratio with the preassembled RNP complexes, and electroporation was carried out using the Neon NxT Transfection System with 100 μL electroporation tips (ThermoFisher, NEON1S). Each pulse was delivered at 1,350 V for 30 ms with a single pulse per sample. The electroporated cell–RNP mixture was immediately transferred into pre-warmed angiogenic media and plated into appropriate culture plates depending on the experimental design.

### PCR and indel analysis by Tracking of Indels by Decomposition (TIDE) and Inference of CRISPR Edits (ICE)

Electroporated cells were cultured until 80-100% confluency in a 12 well plate. DNA was isolated using Quick-Extract DNA Extraction Solution (Lucigen, QE09050) following the manufacturers protocol. PCR was performed using 100 ng of genomic DNA with primers flanking the sgRNA target site, utilizing KAPA HiFi HotStart ReadyMix (Roche, KK2602) according to the manufacturer’s instructions. PCR products were purified with the QIAquick PCR Purification Kit (Qiagen, 28106) and submitted for Sanger sequencing (Genewiz). Primer sequences for PCR amplification and sequencing of the PCR products are shown in **Table 2**. Sequence data from control and edited samples were analyzed using the ICE webtool provided by Synthego (https://ice.editco.bio/#/) or the TIDE webtool (https://tide.nki.nl/#about) to quantify insertion and deletion (indel) frequencies.

**Table 2.**
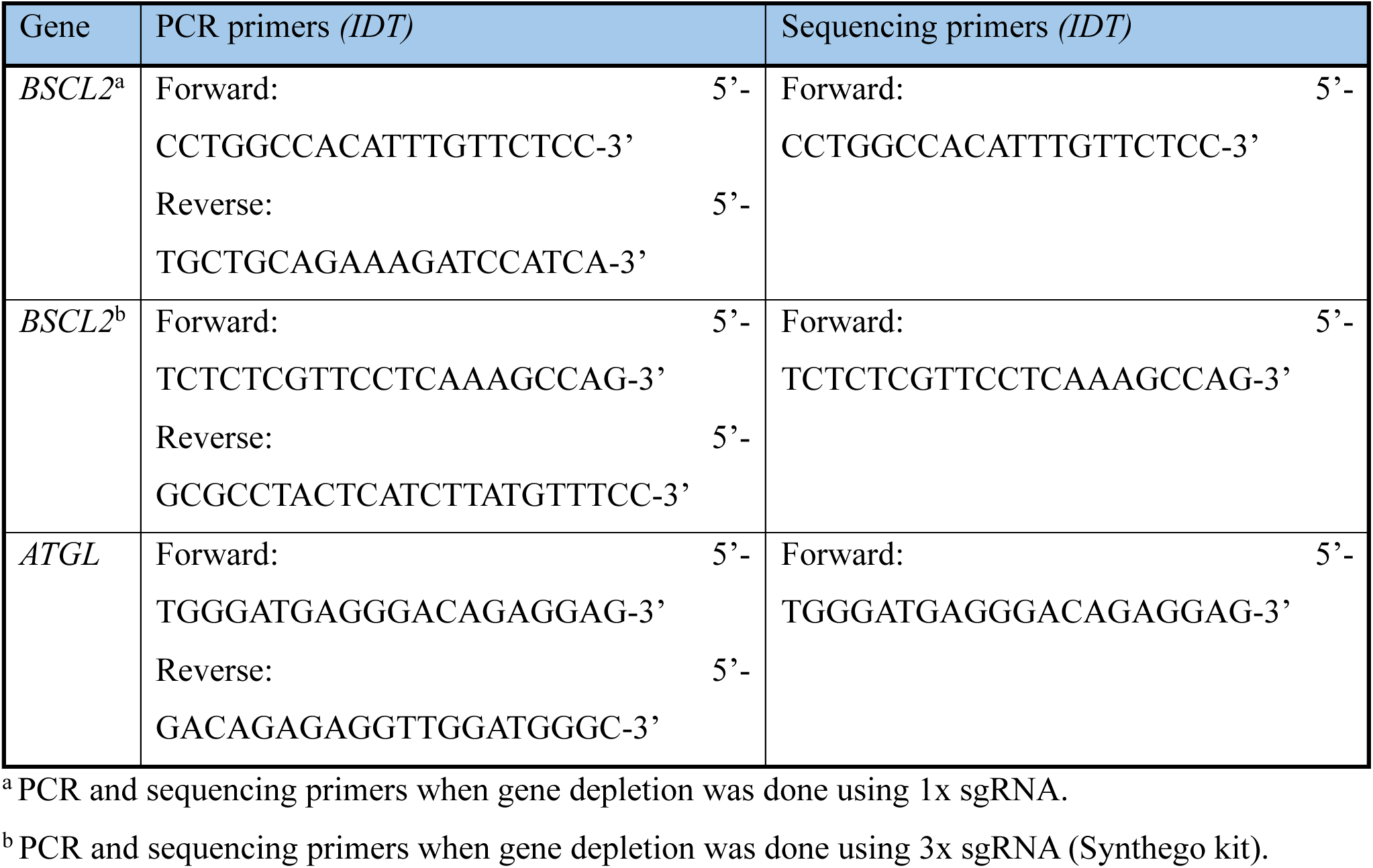
PCR Primer Sequences and sequencing primers of the sgRNA target site.

### Lipid droplet analysis

Lipid droplet analysis was performed using ImageJ software on bright field cell culture images acquired at 10x magnification, with five fields captured per well, as previously described [42]. Image analysis included background subtraction, local contrast enhancement, and particle analysis based on specific circularity parameters for the identification of lipid droplets. This process yielded measurements of lipid droplet area per condition. Statistical analysis of particle area data was conducted using R software.

### Immunofluorescence staining

Cells were cultured on glass coverslips in 24-well plates and fixed with 4% v/v formaldehyde (Thermo Scientific, 28908) in PBS at RT for 15 min with gentle agitation. Cells were permeabilized and blocked in permeabilization buffer containing 1% v/v FBS (Genesee Scientific) and 0.01% w/v digitonin (Sigma, D141) in PBS for 30 min at RT with gentle agitation. Primary antibody against Perilipin-1 (1:150 dilution, Abcam, 61682 and 1:100 dilution, Cell Signaling, 9349), Perilipin 2 (1:400 dilution, ProteinTech, 15294-1-AP) or Seipin (1:100 dilution, Abcam, 106793) was diluted in permeabilization buffer and incubated with the cells for 2 hours at RT with gentle shaking. Following primary antibody incubation, cells were washed with permeabilization buffer. Secondary antibody conjugated to Alexa Fluor 488 (Life Technologies, A11055) was diluted 1:2000 in permeabilization buffer and incubated with cells for 30 min at RT in the dark, followed by washes. Nuclei and lipid droplets were counterstained simultaneously with Hoechst 33342 (1:5000 dilution, Life Technologies, H3570) and LipidTOX Deep Red Neutral Lipid Stain (1:200 dilution, Invitrogen^TM^, H34477) in PBS for 30 min at RT under gentle agitation. Cells were then washed with PBS. Coverslips were mounted onto microscope slides using ProLong™ Gold Antifade Mountant (Thermo Fisher, P36930) and left to cure overnight at RT in the dark. Slides were imaged using the Zeiss Axio Observer Z1 Fluorescence microscope and the Nikon ECLIPSE Ts2R-FL (Diascopic and Epi-fluorescence illumination model) microscope. Exposure times were kept consistent across all samples for each fluorescent dye. At least five images per coverslip were captured. Image analysis was performed using ImageJ software.

For high resolution images, wide-field images were captured on an Axiocam 705 camera using a Zeiss Plan-Apochromat 63x/1.40 Oil DIC M27 objctive a at 250 nm intervals, and deconvolved using 40-60 iterations of the Richardson-Lucy algorithm through the DeconvolutionLab2 ImageJ plugin [43].

### Electron Microscopy

Cells were washed twice with fresh DMEM media (GIBCO/Thermo Fisher, 11995-073) and submitted to the Core Electron Microscopy Facility of University of Massachusetts Chan Medical School for specimen preparation and microscopy.

### Viral infection

Adeno-associated virus serotype 6 (AAV6) vectors encoding either Green Fluorescent Protein (*GFP*) or the *BSCL2* gene were developed by the Horae Gene Therapy Center at the University of Massachusetts Chan Medical School. MSCs were seeded at 60,000cells/cm^2^ and infected upon reaching confluency. Viral particles were resuspended in angiogenic media at a multiplicity of infection (MOI) of 10^5^ and applied directly to the cells. For the removal of the viral particles, full media change into differentiation media was followed after 24h of infection.

### Glycerol and Free Fatty Acid assays

To assess lipolytic activity, cell culture media were collected at designated time points and stored at −80 °C until further use. Glycerol release was measured using the Free Glycerol Reagent (Sigma-Aldrich, F6428) and glycerol standards (Sigma-Aldrich, G7793), following the manufacturer’s protocol. Free fatty acid concentrations were determined using the Free Fatty Acid Assay Kit (Sigma-Aldrich, MAK466), also according to the manufacturer’s instructions. All samples were analyzed without dilution, and each experimental condition was evaluated in technical duplicates.

### Lipid Droplet Isolation

Differentiated adipocytes were lifted from cell cultures using Accutase (GIBCO/Thermo Fisher, A11105-01), washed with PBS, and resuspended in a cracking buffer [50mM HEPES (Sigma, 54457), 3mM MgCl2 (Sigma, M8266), 62.5mM Sucrose (Sigma, S0389), 1x Halt Protease & Phosphatase Inhibitor Cocktail (Thermo Scientific, 1861284) , 0.01 % w/v Digitonin (Sigma, D141)] for single cell suspension. Cells were then passed 5 times through an isobiotec cell homogenizer (Balch Homogenizer) equipped with a 16μm Tungsten Ball clearance to disrupt the plasma membrane. Sucrose concentration was brought to 30% w/v and the sample was layered on top of 45% w/v sucrose cushion, and spun at 4,000 rcf for 30min at 15°C. Floating lipid droplets were collected, weighed, and resuspended in 600μL mass spectrometry grade 100% v/v IPA. The resuspended sample in IPA was vortexed continuously for 15 min at 4°C and spun at 15,000 rcf for 10 min at 4°C. Supernatant was collected and kept in ice. Pellet was resuspended in 600μL IPA, vortexed a second time for 15 min at 4°C, and spun at 15,000 rcf for 10 min at 4°C. Supernatants from both steps were combined, dried down using a SpeedVac at 5°C and submitted to UMass Chan Metabolomics Core for lipidomic analysis as described below. Protein pellet was submitted to UMass Chan Mass Spectrometry Facility for proteomic analysis as described below.

### Lipidomic analysis

Dried samples were reconstituted in 1:1 acetonitrile:isopropanol and loaded into individual LC-MS vials. Lipids were examined using a Vanquish liquid chromatography (LC) platform coupled to a Orbitrap IQ-X Tribrid mass spectrometer equipped with a heated electrospray ionization (HESI) probe (Thermo Electron). LC separation was performed with an Accucore C30 column (2.1 mm x 150 mm, 2.6 µm particle size; ThermoFisher Scientific) using a gradient of solvent A (10 mM ammonium formate, 0.1% formic acid in 60:40 acetonitrile:water) and solvent B (10 mM ammonium formate, 0.1 % formic acid in 88:10:2 isopropanol: acetonitrile: water). Flow rate was 260 µL/min. The LC gradient was: -3min, 30% B; 0 min, 30% B; 2 min, 43% B; 2.1 min, 55% B; 12 min, 65% B; 18.0 min, 85% B; 20 min, 100% B; 25 min, 100% B; 25.1 min, 30% B; 28 min, 30% B. Column temperature was set to 45°C. The autosampler temperature was 15°C, and the injection volume was 2 µL. Source settings were 3800V and 2300V for positive and negative mode, respectively; sheath gas was set to 40 Arb, Aux gas 8 Arb, sweep gas 1 Arb and the ion transfer tube and vaporizer temperature were set to 300°C and 225°C, respectively. Mass spectra were acquired on each sample in positive and negative ionization mode in full-scan mode. Data-dependent MS/MS was run on sample pools using AquireX software (ThermoFisher Scientific) with an exclusion list, inclusion list, and 5 iterative ID injections per ionization mode. For the full-scan acquisition, the resolution was set to 120,000, RF lens was 60%, the automatic gain control (AGC) target was 2x105, the maximum injection time was 100 msec, and the scan range was m/z 200-1500 m/z. For data-dependent MS/MS using AcquireX, the ions in each full scan were isolated in the quadrupole with a 1.5m/z window, fragmented with a normalized HCD stepped collision energy of 20, 30, 40 units, and analyzed at a resolution of 30,000 with an AGC target of 5x104 and a maximum injection time of 54 msec. MSn branching was incorporated with a targeted MS2 trigger for phosphocholine using 32% CID, and a targeted MS3 neutral loss trigger for fatty acids using 35% CID to improve annotation of triglycerides both using the same isolation window, AGC target, and injection time previously specified. The selection of the ions was filtered with an intensity filter of 2x104, a dynamic exclusion window of 2.0 sec, an exclusion list of background ions based on an extraction blank, and an inclusion list. External calibration was performed every 7 days.

Relative quantification of lipids was performed in LipidSearch 5.1 (ThermoFisher Scientific). For “Product Search” the following parameters were used: Precursor tolerance was 5.0 ppm, Product tolerance was 8.0 ppm, Product threshold was set to relative with a value of 1, with no ID range or precursor selection. The “general lc-ms product database” was used and TG and D5labeledTG were selected. The following manual filter was applied in the search: “Rank = 1 and Grade <> “D”. For the Alignment, the following parameters were used: Search type was set to “Product (LCMS)”; Calc. method was “Mean”; Filter type was “Setting filter”, group type was “Data”, RT Tol was 0.15 min, RT Correction Tol was 0.5 min, and S/N Threshold was 3. Post alignment the following manual filters were applied (see Supplemental Table X). Raw peak areas from annotated lipids were exported to excel and filtered using pre-determined quality control criteria: lipids with a background of less than 5x the average abundance of the pooled samples compared with the average abundance of the extraction blanks were removed; lipids with CVs (standard deviation/ mean peak area across triplicate injections of a represented (pooled) biological sample) greater than 20% were removed; lipids with an R value (linear correlation across a four-point dilution series of the representative (pooled) biological sample) less than 0.9 were removed.

### Proteomic analysis

After protein digestion was performed using the S-Trap Micro Spin protocol (PROTIFI) and then dried using a Speed Vac. After drying, the samples were reconstituted in 20 µL of 0.1% formic acid (FA) in 5% acetonitrile (ACN), followed by centrifugation at 16,000 rcf for 16 minutes. Subsequently, 18 µL of the sample was transferred to non-binding HPLC vials and diluted 5 x before injection. 1 µl per sample was injected in triplicate on a TimsTOF Pro2 (Bruker) mass spectrometer, which was coupled to a nanoElute LC system from Bruker. Peptides were then loaded and separated on an in-house-made 75 μm I.D. fused silica analytical column packed with 25-cm ReproSil-Pur C18-AQ (Dr. Maisch, GmbH, 120 Å, 3 μm) particles to a gravity-pulled tip. A 30-minute gradient was employed for data-independent acquisition (DIA)-PASEF, with a flow rate set at 500 nL/min. The captive nano-electrospray voltage was maintained at 1600V, using one column configuration (no trap) and a solvent composition of Solvent A: 0.1% FA in water and Solvent B: 0.1% FA in ACN.

Raw data files (.d format) were processed using Spectronaut version 20.2 (Biognosys) with the Pulsar search engine in directDIA mode (library-free). The analysis employed a two-pass search strategy, beginning with a calibration search at 1% false discovery rate (FDR) to optimize mass accuracy, followed by a comprehensive main search. Precursor and peptide identifications were filtered at 1% FDR at the protein group level using a target-decoy approach. Protein quantification was performed using the MaxLFQ algorithm with normalization applied across all runs. The dataset comprised 18 LC-MS/MS runs across six biological conditions (B1, B2, B3, C1, C2, C3) with three technical replicates per condition. The final analysis identified 182,724 precursors, corresponding to 125,671 unique peptides and 7,645 protein groups at 1% FDR. Post-translational modification (PTM) site localization and stoichiometry calculations were performed, and coefficients of variation (CV) were calculated for quality assessment.

### Bulk RNA-sequencing

Cells were washed with PBS and lysed in TRIzol reagent (ThermoFisher, 15596018). Lysates were transferred to collection tubes and homogenized using the Tissuelyser II (Qiagen). Chloroform (Sigma, C2432) was added at a 1:5 ratio (v/v), and phase separation was performed by centrifugation in phase lock gel heavy tubes. The aqueous phase was collected and mixed with an equal volume of 100% v/v isopropanol (Sigma, I9516), followed by overnight precipitation at −20 °C. RNA was pelleted by centrifugation (12,000 rcf, 15 min, 4°C), washed with 80% v/v ethanol (Fisher Scientific, BP2818500), and resuspended in nuclease-free water. RNA concentration and purity were assessed using a NanoDrop 2000 spectrophotometer (Thermo Scientific).

Library preparation was performed using TruSeq Stranded mRNA Low Throughput Sample Prep Kit (Cat# 20020594, Illumina) according to manufacturer’s instruction. The libraries were sequenced on the NextSeq 500 system (Illumina) using the NextSeq 500/550 High Output Kits v2 (75 cycles; single-end sequencing; Cat# FC-404-2005, Illumina).

Raw paired-end RNA sequencing reads were processed using a custom Bash pipeline executed within a Conda-controlled environment on a Linux high-performance computing cluster. Raw reads were first assessed for quality using FastQC (v0.12.1;[44]). Adapter sequences and low-quality bases were removed using Trimmomatic (v0.39;[45]) with the parameters ILLUMINACLIP:TruSeq3-PE.fa:2:30:10 LEADING:3 TRAILING:3 SLIDINGWINDOW:4:15 MINLEN:36. Post-trimming read quality was re-evaluated with FastQC to confirm adapter removal and quality improvement. Trimmed reads were aligned and quantified using RSEM (v1.3.3;[46]) with the --star option to enable alignment through the STAR aligner (v2.7.11a;[47]). The human reference genome (GRCh38/hg38, Ensembl release 49) was used for read alignment and transcript quantification.

The full pipeline, including all commands and parameters, is available upon request and is reproducible on any Unix-based system with the specified software dependencies. Differentially expressed genes were identified using DESeq2 [48]. Pathway analysis was performed using ToppGene Suite for gene list enrichment analysis and candidate gene prioritization [49].

### Pharmacological PPARγ activation

Cells were treated with 1 μM rosiglitazone (Sigma, R2408) in DMEM following completion of MDI treatment on day 3 of differentiation. Media was replced every 48 hours in the continued presence of rosiglitazone until day 14.

### Adiponectin ELISA

Human adiponectin levels were measured using the Adiponectin Human ELISA Kit (Abcam, ab99968), following the manufacturer’s instructions. Cell culture media samples were diluted 1:100, while serum samples were diluted 1:10 prior to analysis. All samples were previously stored at −80 °C and each condition was evaluated in technical duplicates.

### Lactate dehydrogenase (LDH) assay

Media was collected every 48 hours, followed by a complete media change. Collected media was diluted 1:20 in storage buffer [200 mM Tris-HCl, pH 7.3 (Invitrogen, 15567-027); 10% v/v glycerol (Fisher, G33-500); 1% w/v BSA (Sigma, A3059)] and stored at −80 °C. On the day of the experiment, samples were thawed once and further diluted 1:100 in storage buffer. LDH activity was then measured using the Promega LDH-Glo™ Cytotoxicity Assay Kit (J2381) according to the manufacturer’s instructions.

### In vivo adipose tissue development

Progenitor cells electroporated with sgCTRL or sgBSCL2 guides were plated in 12 15 cm culture plates, grown to confluence and induced as described above. 14 days after induction of differentiation cells were washed twice with PBS and detached using 1x Trypsin-EDTA and 1 mg/mL collagenase I (Worthington-Biochem, LS004196) in PBS. The cells were incubated at 37 °C for 7–10 min and mechanically dissociated by pipetting. Trypsin activity was neutralized using media containing FBS (Genesee Scientific, 25-514), and cells (pelleted and foating) were collected after centrifugation at 500 rcf for 10 min at RT. Cells were resuspended 1:1 in Matrigel (Corning, 356231) and 0.5 ml of cells suspension (containing approximately 5 x 10^6 cells) was injected subcutaneously into the right or left flanks of 6 female and 6 male NSG immunodeficient mice per condition. Blood was collected by cheek bleeding for human adiponectin quantification.

## Data availability

All raw RNA-seq count data, proteomic, and lipidomic datasets generated during this study will be deposited in public repositories upon publication. RNA-seq data will be available through the Gene Expression Omnibus (GEO), proteomic data through the PRIDE repository, and lipidomic data through MetaboLights. All code used for data processing and analysis will be made available via GitHub. Any additional data supporting the findings of this study are available from the corresponding author upon reasonable request.

## ACKNOWLEDGEMENTS

RO1 DK123028 and RO1 DK137403 to SC, a pilot grant from the Li Weibo Institute for Rare Diseases Research to DZ, and fellowships from the Onassis Foundation and the Bodossaki Foundation to IS.

